# Regenerative MRL/MpJ Tendon Cells Exhibit Sex Differences in Morphology, Proliferation, Mechanosensitivity, and Cell-Matrix Remodeling

**DOI:** 10.1101/2022.09.13.507820

**Authors:** Jason C. Marvin, Molly E. Brakewood, Mong Lung Steve Poon, Nelly Andarawis-Puri

## Abstract

Clinical and animal studies have reported the influence of sex on the incidence and progression of tendinopathy, which results in disparate structural and biomechanical outcomes. However, there remains a paucity in our understanding of the sex-specific biological mechanisms underlying effective tendon healing. To overcome this hurdle, our group has investigated the impact of sex on tendon regeneration using the super-healer Murphy Roths Large (MRL/MpJ) mouse strain. Despite a shared scarless healing capacity, we have shown that MRL/MpJ patellar tendons exhibit sexually dimorphic regulation of gene expression for pathways involved in fibrosis, cell migration, and extracellular matrix (ECM) remodeling following an acute midsubstance injury. Moreover, we previously found decreased matrix metalloproteinase-2 (MMP-2) activity in female MRL/MpJ tendons after injury. Thus, we hypothesized that MRL/MpJ scarless tendon healing is mediated by sex-specific and temporally distinct orchestration of cell-ECM interactions. Accordingly, the present study comparatively evaluated MRL/MpJ tendon cells under two-dimensional (glass) and three-dimensional (nanofiber scaffolds) culture platforms to examine cell behavior under biochemical and biophysical cues associated with tendon homeostasis and healing. Female MRL/MpJ cells showed reduced 2D migration and spreading area accompanied with enhanced mechanosensing, 2D ECM alignment, and fibronectin-dependent cell proliferation. Interestingly, female MRL/MpJ cells cultured on 3D isotropic scaffolds showed diminished ECM deposition and alignment. Regardless of culture condition and sex, MRL/MpJ cells outperformed B6 cells and elicited a universal regenerative cellular phenotype. These results illustrate the utility of these *in vitro* systems for elucidating regenerative tendon cell biology, which will facilitate the long-term development of more equitable therapeutics.

## 1. Introduction

Tendons are dense connective tissues that enable skeletal locomotion and joint stability by transmitting tensile forces from muscle to bone. The tendon extracellular matrix (ECM) primarily consists of highly aligned, hierarchically organized type I collagen fibers, water, and non-collagenous components (e.g., proteoglycans, elastin, tenascin-C, and other glycoproteins) that impart the dynamic loading capabilities of the tissue.^1–3^ Due to their relatively low vascularity and cellularity, tendon ruptures in adult mammals heal poorly with the formation of disorganized scar tissue (‘scar-mediated healing’) with compromised biomechanical integrity.^4–6^ The treatment and management of tendon and ligament injuries is further complicated in that they display sex differences in structure and mechanical properties during growth^7–10^, adaptation and collagen synthesis in response to physiological exercise^11,12^, and post-operative recovery outcomes in both patients and large animal models.^13–17^ Given the high clinical prevalence of tendinopathies, there is a critical need to address a paucity in our fundamental knowledge of sex-specific biological mechanisms underlying effective tendon healing.

To expand our understanding of the cellular and molecular mechanisms underpinning tendon regeneration, several vertebrate animal models have been explored, including amphibians, zebrafish^18^, neonatal mice^19^, and select transgenic strains such as the super-healer Murphy Roths Large (MRL/MpJ) mouse.^20–25^ In particular, our group and others have established that MRL/MpJ mice elicit a scarless healing capacity following acute injury of the patellar, flexor digitorum longus, and supraspinatus tendons compared to those of non-healer C57BL/6 (B6) mice.^20–23,25,26^ We have previously determined using *ex vivo* organ culture that the local tissue environment rather than systemic contributions (i.e., the immune response) serves as the primary driver of MRL/MpJ tendon healing^23^. We have also demonstrated that the administration of decellularized MRL/MpJ tendon injured/provisional ECM and cell-derived soluble factors *in vitro* shifts B6 cells toward a regenerative cellular phenotype^23,27^, which illustrates the promising utility of isolating constituents of the MRL/MpJ tendon biological environment to guide the development of mechanistically-informed therapeutics. Furthermore, we have shown sexually dimorphic regulation of wound healing-associated genes and matrix metalloproteinase-2 (MMP-2) activity during MRL/MpJ acute tendon healing^24^. Together, these findings raised an important question of whether these unique genetic profiles and temporal shifts in ECM remodeling processes reflect sex differences in MRL/MpJ cell behavior, thereby suggesting potentially different biological and cell signaling pathways that are necessary to achieve similar regenerative healing outcomes.

In the present study, we comparatively evaluated MRL/MpJ cell behavior using *in vitro* metrics^23,27^ known to differ between male B6 and MRL/MpJ cells consistent with scarless healing *in vivo*. Our primary objective was to identify sex differences in MRL/MpJ cells that may be integral in distinctively modulating the early healing cascade towards the improved structural alignment characteristic of MRL/MpJ tendon healing. To assess how changes in ECM and topographical cues impact these cellular functions, we cultured primary mouse patellar tendon cells on two-dimensional (2D; glass) substrates and three-dimensional (3D; commercial nanofiber scaffolds) platforms with tightly controlled fiber architectures representative of healthy and injured tendon environments. Using these 2D and 3D systems, we analyzed single-cell and confluent cell features, including cytoskeletal organization, proliferation, migration, mechanosensitivity, and ECM deposition. We hypothesized that (i) female MRL/MpJ cells would exhibit comparable morphological features but exhibit decreased cell migration and increased ECM deposition due to decreased catabolic activity compared to male MRL/MpJ tendon cells, and (ii) sex differences in MRL/MpJ cell behavior would become more pronounced under 3D culture conditions.

## 2. Methods

### 2.1. Primary tendon cell culture, reagents, and 3D nanofiber scaffolds

B6 and MRL/MpJ mice at 16-to-18 weeks of age were bred in-house with the original dame and sire purchased from The Jackson Laboratory (B6: #000664, MRL/MpJ: 000486). Mice were housed with up to five animals per cage in an alternating 12-hour light/dark cycles with ab libitum access to chow and water. Under Cornell University Institutional Animal Care and Use Committee (IACUC) approval, patellar tendon cells were isolated from male B6 and male and female MRL/MpJ mice as described previously.^23^ After animals were euthanized, patellar tendons from both limbs of five animals were harvested, removed of the fat pad and sheath, and digested together with a combination of collagenase I (2 mg/mL) and collagenase type IV (1 mg/mL) in serum-free DMEM for up to 2 hours at 37°C on a rocking shaker. Digest solutions were passed through a 70-μm strainer to seed single-cell suspensions (passage 0).

Cells were cultured in basal media consisting of low-glucose Dulbecco’s modified Eagle’s medium (DMEM; Corning, #10-014) containing 10% (v/v) lot-selected fetal bovine serum (FBS; Corning, #35-015-CV) and 1% (v/v) penicillin-streptomycin-amphotericin B (PSA; Thermo Fisher Scientific, #15240062). All experiments were performed at passage 3 with media replaced every 2 days. Cells used in this study were negative for mycoplasma (Lonza, #LT07-703). To recapitulate compositional cues during tendon homeostasis and injury, ECM-coated substrates with rat tail collagen I (Thermo Fisher Scientific, #A1048301) or human plasma fibronectin (#33-016-015), respectively, were added at a concentration of 50 μg/mL overnight at 4°C. Coatings were rinsed twice with 1X phosphate-buffered saline (PBS) immediately prior to use.

For 2D culture experiments, cells were cultured on glass-bottom 96-well plates (Greiner Bio-One, #655892). For 3D culture experiments, cells were cultured on commercial polycaprolactone (PCL) nanofiber scaffolds (20-μm thickness) with aligned (NanoAligned; Nanofiber Solutions, #9602) and disorganized (NanoECM; Nanofiber Solutions, #9601) fiber topographies that mimic healthy and injured tendon structure, respectively, as per the manufacturer’s instructions (**Fig. 1**). Additionally, for 3D culture experiments, male B6 tendon cells served as a control to comparatively evaluate previously established *in vitro* metrics that were different between male B6 and MRL/MpJ tendon cells.^23,27^

**Figure 1.**
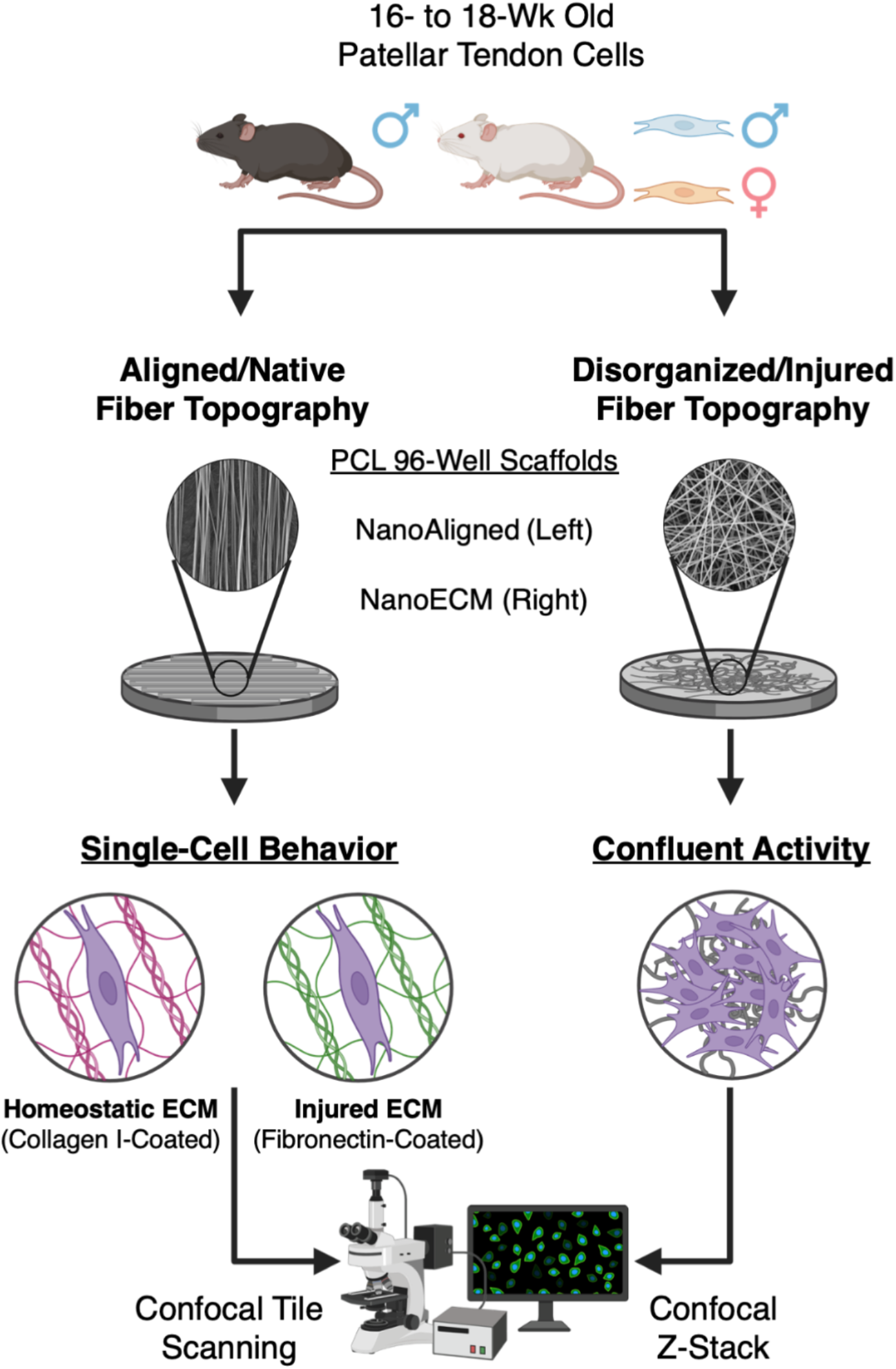
Experimental schematic for the comparative evaluation of sex differences in MRL/MpJ tendon cell behavior using 3D nanofiber scaffolds *in vitro*. For 3D culture experiments, tendon cells were cultured on ECM-coated NanoAligned and uncoated NanoECM scaffolds with anisotropic/aligned and isotropic/disorganized fiber topographies, respectively. Male B6 cells served as a control to confirm whether previously established *in vitro* metrics of the MRL/MpJ cellular phenotype are consistent between 2D and 3D culture conditions. Singlecell behavior and confluent cell activity was assessed using immunofluorescence staining and confocal imaging of cells cultured directly within the nanofiber scaffolds.

### 2.2. Immunofluorescence (IF) staining and confocal microscopy

Cells were fixed with 2% paraformaldehyde (PFA) in 1X PBS for 10 minutes followed by 4% PFA for 20 minutes at room temperature. All subsequent immunostaining steps were conducted on an orbital shaker at 50 RPM. Cells were then washed thrice with 1X PBS for 5 minutes per wash, permeabilized with 0.1% Triton X-100 in 1X PBS for 15 minutes, and blocked with 3% bovine serum albumin (BSA) in 1X PBS for 1 hour at room temperature. Primary antibodies in 3% BSA were incubated as specified (**Table 4.1**). After the primary incubation, cells were washed thrice with 1X PBS for 5 minutes per wash before being incubated with Alexa Fluor secondary antibodies in 3% BSA for 1 hour at room temperature while protected from light. Lastly, cells were washed thrice with 1X PBS for 5 minutes per wash, counterstained with 1 μg/mL of 4′,6-diamidino-2-phenylindole (DAPI; Thermo Fisher Scientific, #D1306) for 30 minutes, and finally rinsed twice and mounted with 1X PBS before imaging. To label extracellular fibronectin, the primary antibody was incubated prior to permeabilization.

**Table 1.** List of antibodies used for immunofluorescence staining.

| Antibody/Reagent | Manufacturer | Catalog Number | Dilution and Use |
| --- | --- | --- | --- |
| Alexa Fluor™ Plus<br>555 Phalloidin | Thermo Fisher<br>Scientific | A30106 | 1:400, 1 hour at room<br>temperature |
| Vinculin | Sigma-Aldrich | V9131 | 1:400, overnight at 4°C |
| YAP | Santa Cruz<br>Biotechnology, Inc. | sc-101199 | 1:200, 2 hours at room<br>temperature |
| Ki-67 | Abcam | ab15580 | 1:1000, overnight at 4°C |
| Fibronectin | Abcam | ab45688 | 1:400, overnight at 4°C |
| Connexin-43 | Abcam | ab11370 | 1:1000, overnight at 4°C |
| Vimentin | Abcam | ab92547 | 1:250, overnight 4°C |

Fluorescent images were acquired on a Zeiss LSM710 confocal microscope (Zeiss) using a 25X or 40X magnification water-immersion objective. For NanoAligned samples, one 2D tile scan (3188 μm by 3188 μm) was taken for each scaffold. For NanoECM samples, five representative full-thickness Z-stacks were taken for each scaffold. Identical acquisition parameters were used across samples for each independent experiment. Image analysis was performed in ImageJ (National Institutes of Health) by a blinded user.

### 2.3. Cellular senescence and migration

To assess 2D cellular senescence, MRL/MpJ cells were seeded at 5,000 cells/well onto uncoated glass substrates in basal media and cultured for 24 hours. Cells were then stained for β-galactosidase (β-gal) using a commercial kit (Cell Signaling Technology, #9860S) and imaged at 4X magnification on an inverted brightfield microscope. The percentage of senescent cells (*N* = 5 per group averaged from 3 wells per *N*) was determined as the number of cells positive for ß-gal divided by the total number of cells.

To measure 2D cell migration, MRL/MpJ cells were seeded at 7,500 cells/well onto collagen I-coated glass substrates in basal media and cultured for 12 hours. Cells were subsequently serum starved in low-serum (1% FBS) media for 18 hours to induce cell cycle synchronization, manually scratched with a 200 μL pipette tip, rinsed once with serum-free media, and then cultured in low-serum media. Transmitted light images were immediately taken (0 hour post-scratch) on a Spectra Max i3X Multi-Mode Microplate Reader (Molecular Devices) to determine the original scratch region. At 24 hours post-scratch, cells were labeled with 2 μM calcein AM for 30 minutes at 37°C and 5% CO_2_. Fluorescent images were then taken on the Spectra Max i3X Multi-Mode Microplate Reader to quantify the total number of cells that migrated into the defect region *(N* = 3 per group averaged from 3 wells per *N*).

### 2.4. Single-cell behavior in 2D and 3D culture

To assess single-cell behavior, MRL/MpJ cells were seeded at 750 cells/well onto collagen I-or fibronectin-coated glass substrates and NanoAligned scaffolds in basal media and cultured for 18 hours. Cell morphology, proliferation, and mechanosensitivity were analyzed (*N* = 54 cells or 72 cells for 2D and 3D culture, respectively) by staining for F-actin, Ki-67, and Yes-associated protein (YAP) and vinculin, respectively. Focal adhesion measurements were limited to 2D culture experiments due to the lack of discrete focal adhesion sites detected on NanoAligned samples. Cell morphology was measured for spreading area, perimeter, aspect ratio (only for 3D culture), and circularity as defined by the following:

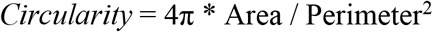

Cellular protrusions were manually counted and traced for all cytoplasmic extensions over 25 μm in length. For NanoAligned samples, the percentage of Ki-67-positive cells (*N* = 8-9 scaffolds per group) was averaged from three randomly selected ROIs per scaffold. Vinculin-positive focal adhesions were quantified on a semi-automated basis as described previously.^28^ YAP was analyzed as the nuclear-to-cytoplasmic mean intensity ratio (nuclear localization).

### 2.5. Confluent cell activity and fibronectin deposition in 2D and 3D culture

To assess confluent cell activity, MRL/MpJ cells were seeded at 2,000 cells/well or 3,500 cells/well on uncoated glass substrates and NanoECM scaffolds, respectively, in basal media and cultured for 7 days. Cytoskeletal organization, intercellular communication, and ECM deposition were analyzed (*N* = 45 images or Z-stacks for 2D and 3D culture, respectively) by staining for F-actin or vimentin, connexin-43 gap junctions, and fibronectin, respectively. To measure cytoskeletal and ECM alignment, five randomly selected elliptical ROIs for each image/Z-stack were quantified using the OrientationJ plugin^29^ in ImageJ and averaged. A coherency value close to 1 indicated anisotropy with uniform alignment whereas a value close to 0 indicated isotropy with no preferential alignment. To measure cellular organization, nuclear orientation was calculated for cells cultured on NanoECM samples. Integrated density measurements for fibronectin and connexin-43 were normalized to the total number of cells per image.

### 2.6. Statistical analysis

All experiments were performed with a minimum of 3 biological replicates (i.e., independent cell isolation batch). Cellular senescence, migration, proliferation, and the percentage of cells with protrusions were compared between sexes using an unpaired Student’s t-test (2D culture) or one-way analysis of variance (ANOVA) with post-hoc Tukey (3D culture). The number of cellular protrusions were compared between sexes using a Kolmogorov-Smirnov test (2D culture) or a Kruskal-Wallis test with post-hoc Dunn’s (3D culture). Integrated density data for fibronectin and connexin-43 were compared between sexes using a Mann-Whitney *U* test (2D culture) or Kruskal-Wallis test with post-hoc Dunn’s (3D culture). All other confluent and single-cell outcome measures were compared between sexes using a Welch’s t-test (2D culture) or one-way Welch ANOVA with post-hoc Games-Howell (3D culture). The Shapiro-Wilk test was performed to assess the normality of our datasets. Statistical analyses were done using GraphPad Prism 9.3.1 (GraphPad Software). Statistical significance was set at *P* < 0.05. Data are shown as mean + standard error of the mean (SEM).

## 3. Results

### 3.1. Female MRL/MpJ cells show comparable senescence but decreased cell migration

No sex differences in cellular senescence were detected for MRL/MpJ cells (**Fig. 2A-B**). There also were no differences in the number of senescent cells or total cell number (data not shown). Supporting our hypothesis, scratch assays (**Fig. 2C**) revealed that significantly less female cells migrated into the defect site compared to male cells after 24 hours (**Fig. 2D**).

**Figure 2.**
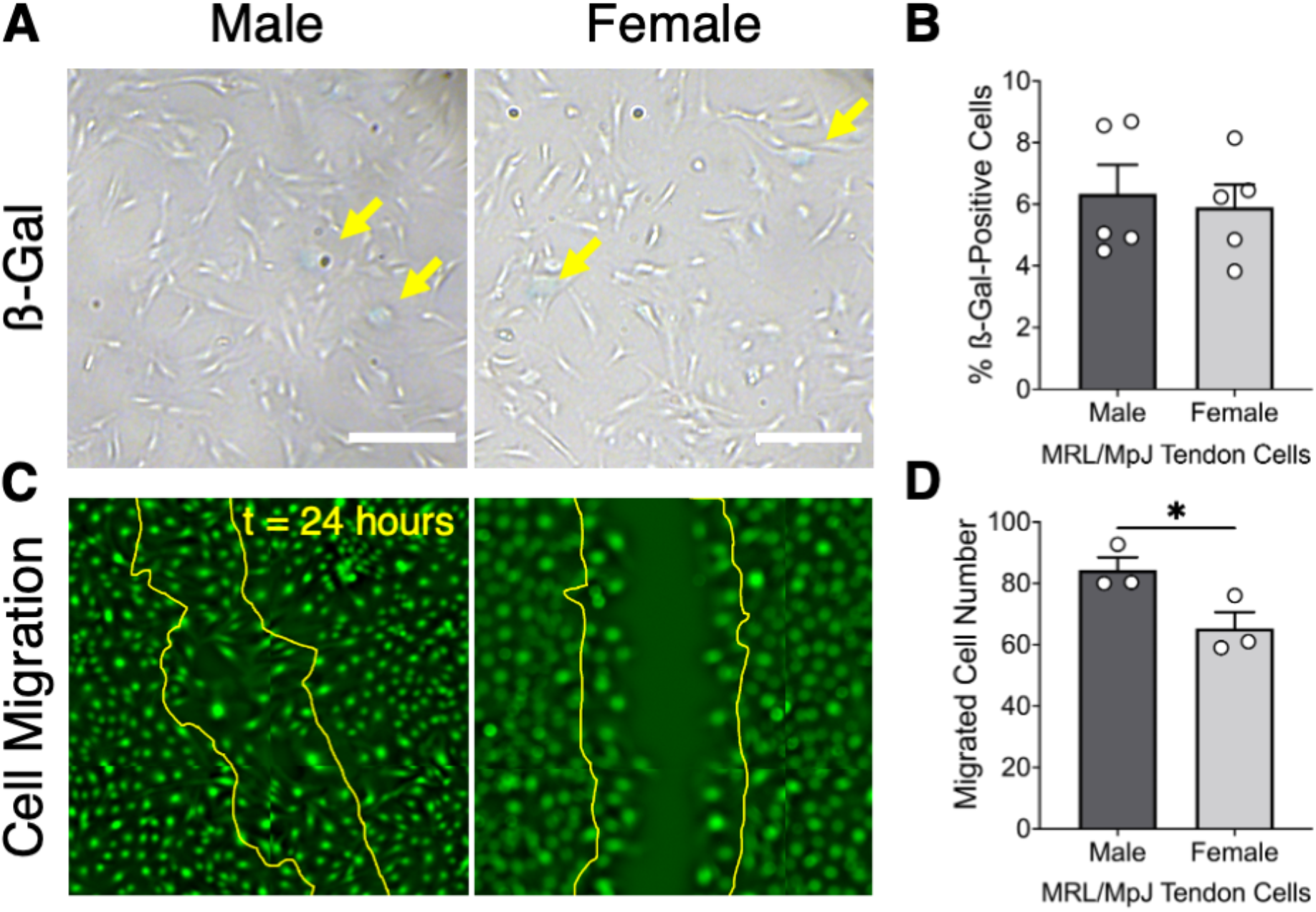
Comparison of 2D cellular senescence and cell migration between male and female MRL/MpJ tendon cells. (**A**). Representative brightfield images of MRL/MpJ cells at a moderate density stained for ß-gal as an indicator of cellular senescence. Yellow arrows indicate ß-gal-positive/senescent cells. Scale bar, 250 μm. (**B**) Quantification of ß-gal staining revealed no sex differences in cellular senescence for MRL/MpJ cells. (**C**) Representative fluorescent images of confluent MRL/MpJ cells cultured on collagen I-coated glass substrates at t = 24 hours after scratching. Yellow outline indicates the original defect at t = 0 hour. (**D**) Significantly less female than male MRL/MpJ cells migrated into the defect site after t = 24 hours. Data is presented as mean + SEM. \**P* < 0.05.

### 3.2. Female MRL/MpJ cells are smaller with enhanced focal adhesions and cellular protrusions

Female cells exhibited significantly less 2D spreading area than male cells regardless of ECM substrate (25.4% and 42.3% less; *P* = 0.047 and 0.021 for collagen I and fibronectin, respectively), in addition to lower circularity (*P* = 0.077) and perimeter (*P* = 0.14) on collagen I and fibronectin, respectively (**Fig. 3A-B**). The number of actin-rich protrusions were comparable between sexes regardless of ECM substrate, but female cells displayed a significantly greater percentage of cells with protrusions (*P* = 0.0075) and total protrusion length (*P* = 0.027) than male cells on fibronectin (**Fig. 3C**). When normalized to cell spreading area to account for sex differences in cell morphology, female cells formed significantly more focal adhesion sites (30.8% and 15.9% higher; *P* < 0.0001 and 0.033 on collagen I and fibronectin, respectively) and coverage (*P* = 0.0076 and 0.029 on collagen I and fibronectin, respectively) than male cells regardless of ECM substrate (**Fig. 3D**), implicating altered cell-ECM adhesion as a potential mediator of these changes in cell migration and morphology.

**Figure 3.**
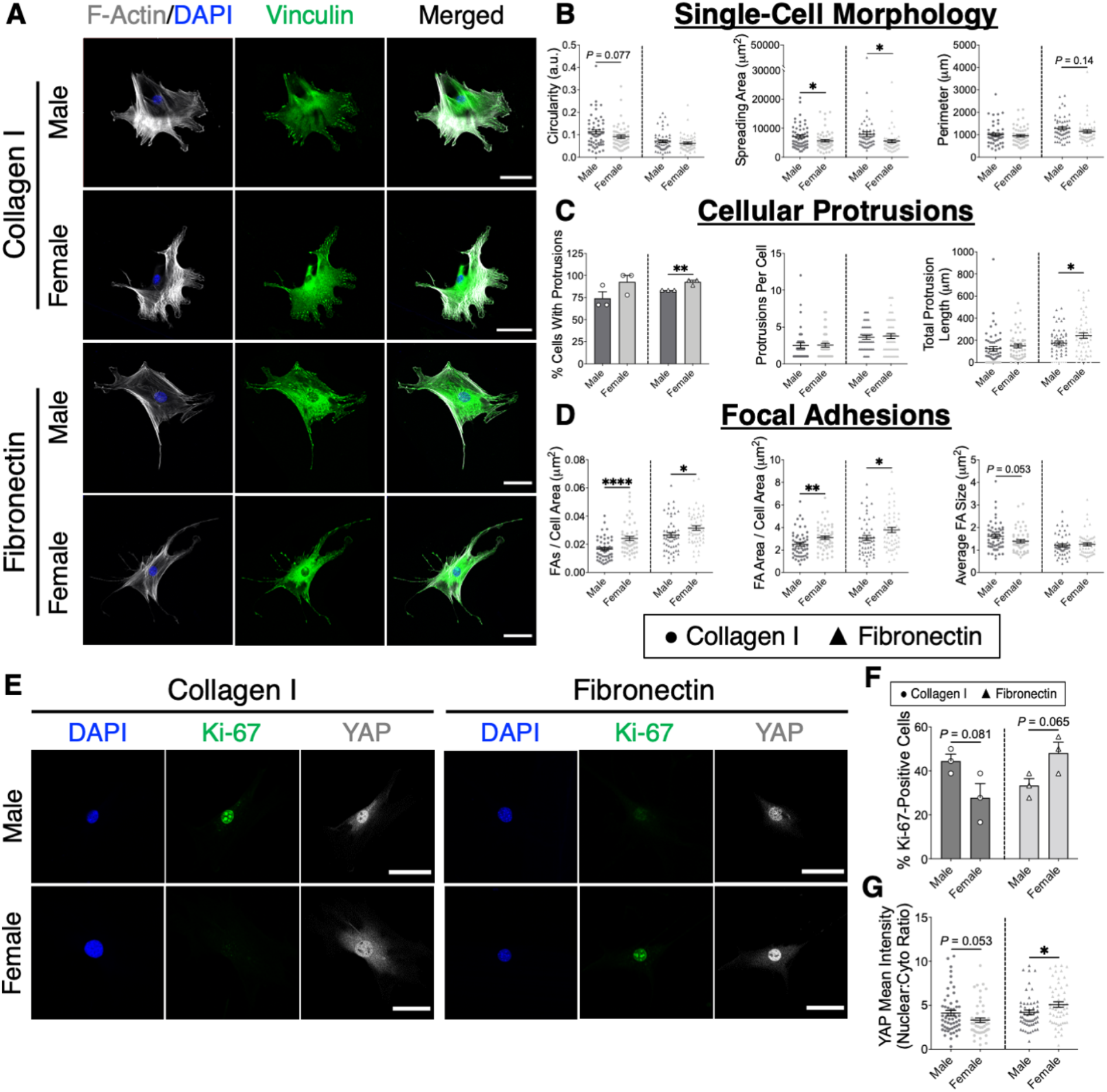
Assessment of sex differences in 2D single-cell morphology, proliferation, and mechanosensitivity in MRL/MpJ tendon cells. (**A**) Representative IF staining of the cytoskeleton (F-actin, gray) and focal adhesions (vinculin, green) for MRL/MpJ tendon cells cultured at a low density on collagen I- and fibronectin-coated glass substrates. Scale bar, 50 μm. (**B-D**) Quantification of cell morphological parameters (**B**), cellular protrusions (**C**), and focal adhesion assembly (**D**). Female MRL/MpJ tendon cells exhibited significantly decreased spreading area and increased focal adhesion formation, as well as enhanced cellular protrusions on fibronectin. (**E**) Representative IF staining of cell proliferation (Ki-67, green) and mechanosensitivity (YAP, gray) for MRL/MpJ tendon cells cultured at low density on collagen I- and fibronectin-coated glass substrates. Scale bar, 50 μm. (**F-G**) Quantification of the percentage of Ki-67-positive staining (**F**) and YAP nuclear localization (**G**) revealed ECM-dependent sex differences in cell proliferation and mechanosensitivity. Circles and triangles indicate individual data points for cells cultured on collagen I and fibronectin, respectively. \**P* < 0.05, \*\**P* < 0.01, \*\*\*\**P* < 0.0001.

Female cells showed less proliferation and YAP nuclear localization than male cells on collagen I (16.66% and 24.0% lower; *P* = 0.081 and 0.053 for Ki-67 and YAP, respectively) (**Fig. 3E**). In contrast, female cells showed greater proliferation and YAP nuclear localization (14.8% and 21.4% higher; *P* = 0.065 and 0.032 for Ki-67 and YAP, respectively) than male cells on fibronectin. Collectively, these findings indicate that female MRL/MpJ cells harbor a heightened activation response to wound healing-associated biochemical cues under 2D culture conditions.

### 3.3. Sex differences in MRL/MpJ cell morphology, proliferation, and mechanosensitivity are preserved in 3D culture on aligned nanofiber scaffolds

To assess how 3D physical confinement reflective of healthy tendon architecture impacts cellular function, we next characterized previously established 2D single-cell metrics for B6 and MRL/MpJ cells on ECM-coated NanoAligned scaffolds with anisotropic fiber alignment *in vitro* (**Fig. 4A-B**). Visually, all cells formed cellular protrusions and oriented longitudinally along the direction of the parallel fibers. Contrary to our previous 2D data, male B6 cells exhibited greater or comparable spreading area, perimeter, and cellular protrusions than MRL/MpJ cells regardless of ECM substrate (**Fig. 4C-D**). However, B6 cells showed significantly lower aspect ratio (i.e., elongation indicative of native morphology) than male (24.3% lower; *P* = 0.018) and female MRL/MpJ cells (36.2% lower; *P* = 0.0013) on collagen I. Consistent with our 2D data for Ki-67 staining (**Fig. S1**), male B6 cells displayed significantly greater proliferation than male MRL/MpJ cells (15.5% higher; *P* = 0.022) on collagen I (**Fig. 4E**). Contrary to our 2D data for YAP in our other ongoing studies, male B6 cells showed significantly greater YAP nuclear localization than male MRL/MpJ cells (16.2% higher; *P* = 0.03) (**Fig. 4E**).

**Figure 4.**
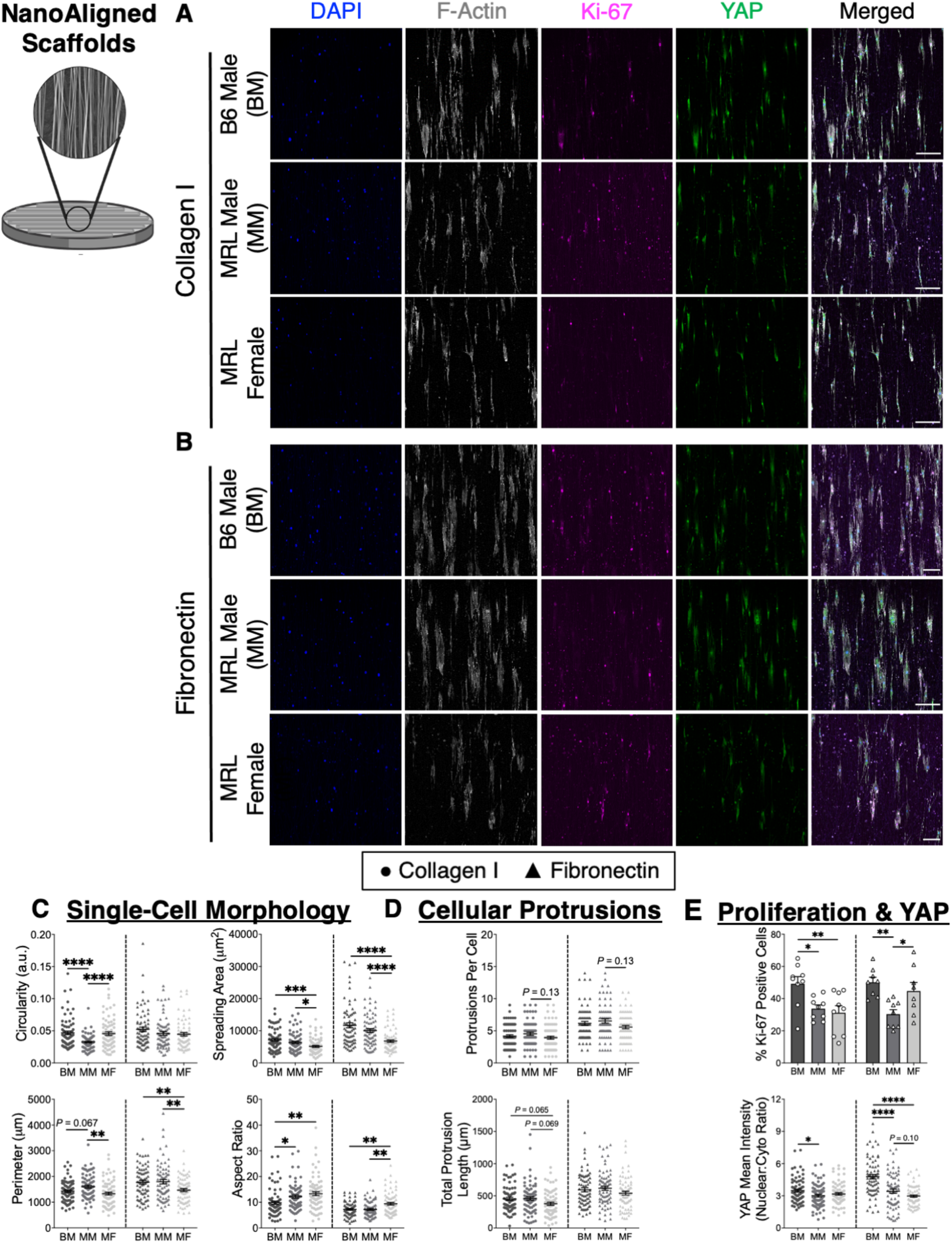
Assessment of 3D single-cell morphology, proliferation, and mechanosensitivity of B6 and MRL/MpJ tendon cells cultured on NanoAligned scaffolds. (**A-B**) Representative IF staining of the cytoskeleton (F-actin, gray), cell proliferation (Ki-67, magenta), and mechanosensitivity (YAP, green) for cells cultured at a low density on collagen I- (**A**) and fibronectin-coated (**B**) NanoAligned scaffolds. Scale bar, 200 μm. (**C-E**) Quantification of cell morphological parameters (**C**), cellular protrusions (**D**), and Ki-67-positive staining and YAP nuclear localization (**E**). Consistent with 2D culture, female MRL/MpJ cells exhibited significantly decreased spreading area compared with male MRL/MpJ cells. On fibronectin-coated scaffolds, female MRL/MpJ cells displayed significantly increased elongation and proliferation in addition to decreased YAP nuclear localization despite comparable activity on collagen-coated scaffolds, indicating a sex-specific response to healing-associated biochemical cues. Circles and triangles indicate individual data points for cells cultured on collagen I and fibronectin, respectively. \**P* < 0.05, \*\**P* < 0.01, \*\*\**P* < 0.001, \*\*\*\**P* < 0.0001.

Consistent with our 2D sex differences data, female MRL/MpJ cells exhibited significantly lower spreading area than male MRL/MpJ cells regardless of ECM substrate (23.4% and 48.4% lower; *P* = 0.028 and < 0.0001 for collagen I and fibronectin, respectively) (**Fig. 4C**). Female MRL/MpJ cells also exhibited significantly lower perimeter than male MRL/MpJ cells regardless of ECM substrate (19.9% and 21.9% lower; *P* = 0.0039 and 0.0066 on collagen I and fibronectin, respectively), which was attributed to male cells forming more protrusions on a per cell basis (*P* = 0.13 for both) (**Fig. 4C-D**). Supporting our hypothesis, female MRL/MpJ cells showed significantly greater aspect ratio than male MRL/MpJ cells on fibronectin (31.6% higher; *P* = 0.0015). Further consistent with our 2D sex differences data, female MRL/MpJ cells displayed significantly greater proliferation than male MRL/MpJ cells on fibronectin (14.4% higher; *P* = 0.036) (**Fig. 4E**). Lastly, female MRL/MpJ cells showed significantly lower YAP nuclear localization than male MRL/MpJ cells on fibronectin (15.4% lower; *P* = 0.1) (**Fig. 4E**).

### 3.4. Confluent female MRL/MpJ cells undergo enhanced matrix remodeling

After 7 days of culture to confluency in full-serum media (**Fig. 5A-B**), female cells showed significantly greater total cell number (20.3% higher; *P* = 0.0068) and ECM alignment (*P* = 0.0087) than male cells (**Fig. 5C and F**). There were no sex differences in fibronectin deposition normalized on a per cell basis and cytoskeletal alignment between sexes, but female cells deposited significantly more total fibronectin than male cells (data not shown) (**Fig. 5D-E**). Female cells exhibited lower connexin-43 expression normalized on a per cell basis than male cells (5.64% lower; *P* = 0.09) (**Fig. 5G**), which was largely attributed to differences in total cell number.

**Figure 5.**
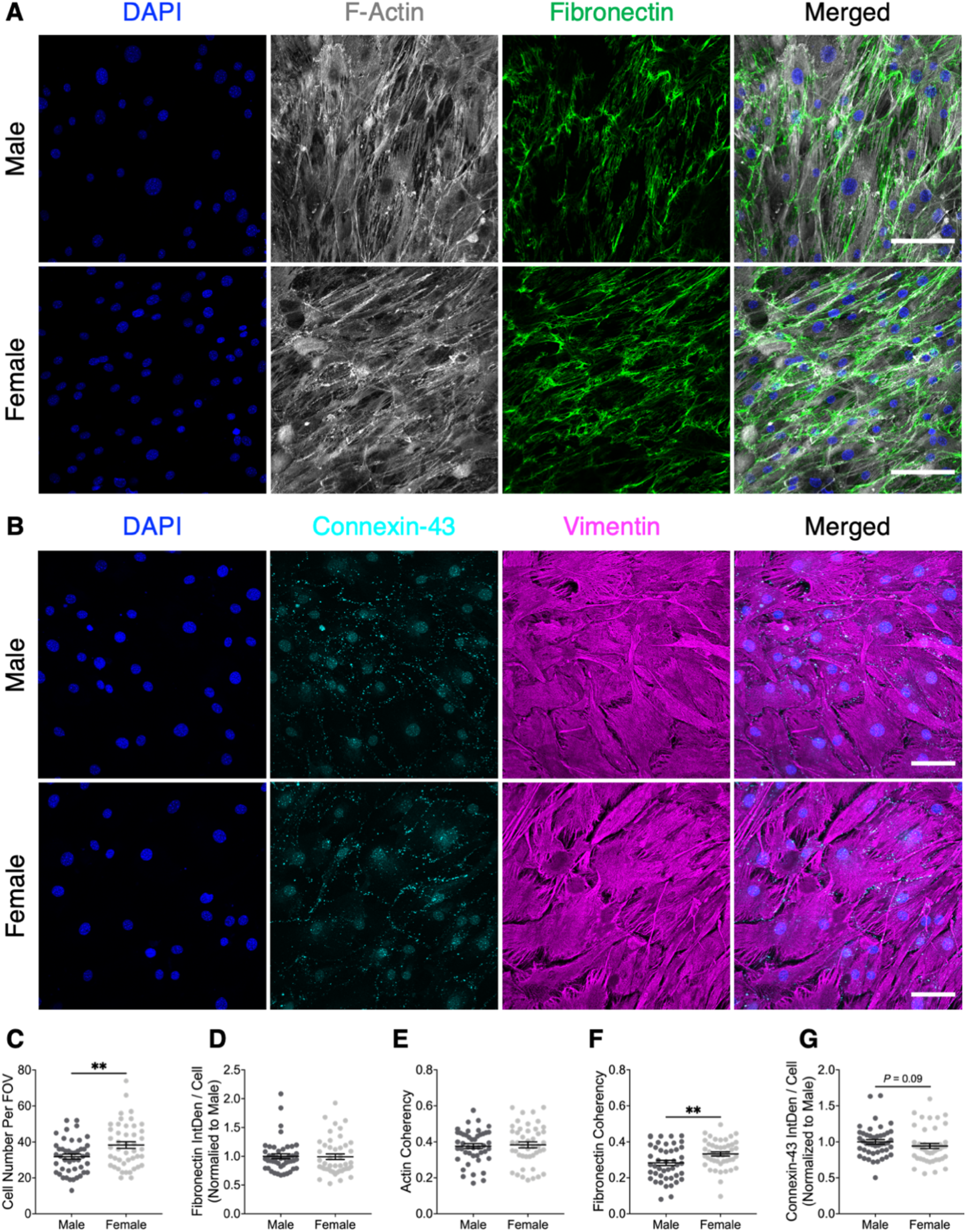
Analysis of 2D sex differences in cytoskeletal and ECM organization, ECM deposition, and gap junction-mediated intercellular communication in MRL/MpJ tendon cells cultured to confluency on uncoated glass substrates. (**A**) Representative IF staining of the cytoskeleton (F-actin, gray) and fibronectin (green) for confluent MRL/MpJ cells. Scale bar, 100 μm. (**B**) Representative IF staining of connexin-43 gap junctions (cyan) and cytoskeleton (vimentin, magenta) for confluent MRL/MpJ cells. Scale bar, 100 μm. (**C-G**) Quantification of cell number (**C**), fibronectin deposition (**D**), cytoskeletal (**E**) and fibronectin alignment (**F**), and intercellular communication via connexin-43 expression (**G**). Female MRL/MpJ cells exhibited significantly increased cell number and fibronectin alignment and decreased intercellular communication at confluency compared to that of male cells. \*\**P* < 0.01.

### 3.5. Cell-ECM remodeling is disrupted in confluent female MRL/MpJ cells under 3D culture on disorganized nanofiber scaffolds

To examine how a disorganized 3D surface topography reflective of the acute tendon injury environment impacts wound healing-associated cell activity, we next evaluated established 2D confluent metrics for B6 and MRL/MpJ cells on uncoated NanoECM scaffolds with isotropic fiber alignment *in vitro* (**Fig. 6A-B**). Contrary to our 2D cell number data at confluency in our other ongoing studies, male B6 cells exhibited a significantly greater total cell number than male MRL/MpJ cells (31.6% higher; *P* < 0.0001) in male B6 cells compared to male MRL/MpJ cells (**Fig. 6C**). However, supporting our hypothesis, male MRL/MpJ cells displayed significantly greater fibronectin deposition (2.04-fold higher, *P* < 0.0001), cytoskeletal (*P* < 0.0001) and ECM alignment (*P* < 0.0001), and connexin-43 expression (1.39-fold higher, *P*<0.0001) than male B6 cells (**Fig. 6D-G**).

**Figure 6.**
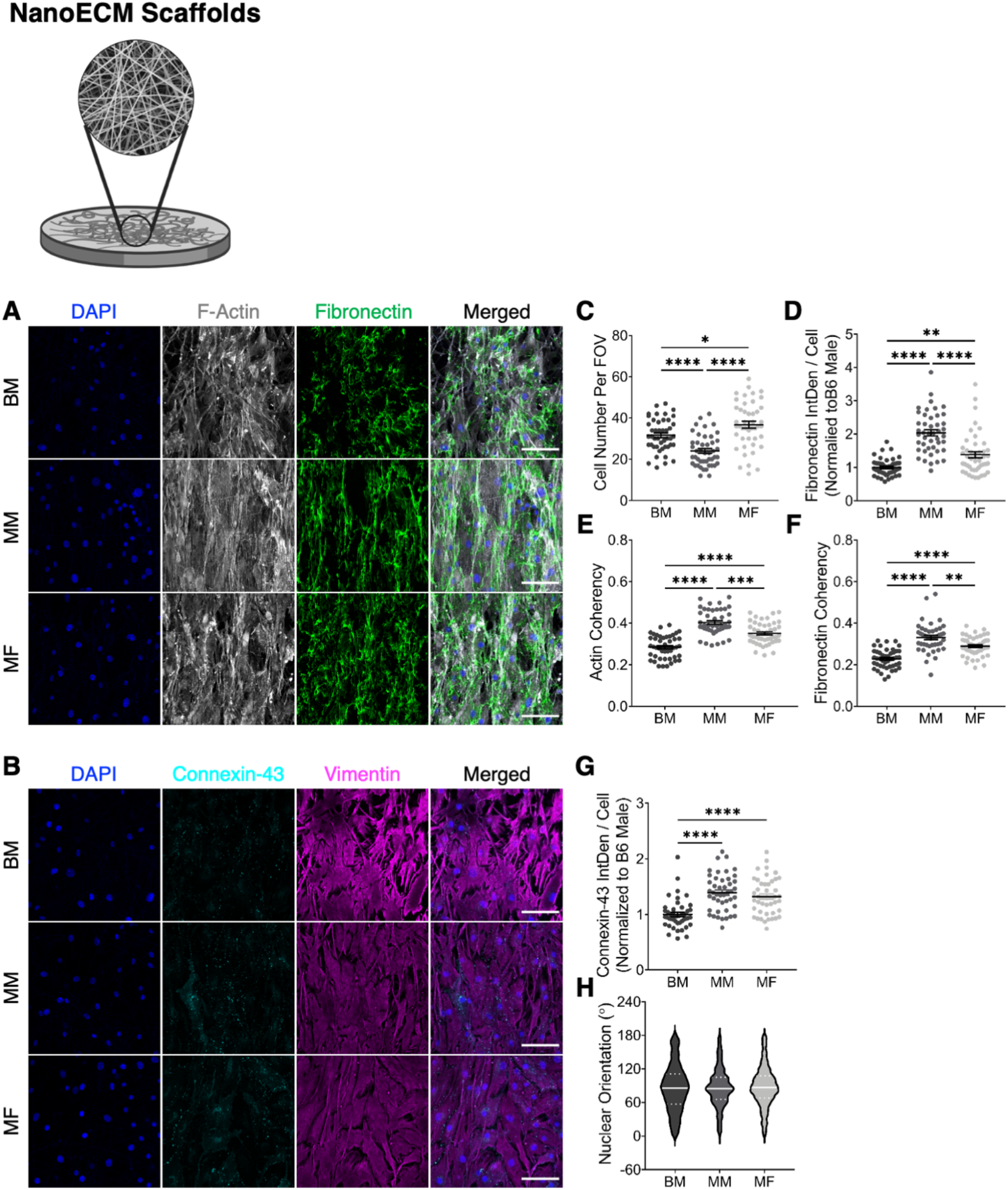
Analysis of 3D sex differences in cytoskeletal and ECM organization, ECM deposition, and gap junction-mediated intercellular communication in MRL/MpJ tendon cells cultured to confluency on uncoated NanoECM scaffolds. (**A**) Representative IF staining of the cytoskeleton (F-actin, gray) and fibronectin (green) for confluent MRL/MpJ cells. Scale bar, 100 μm. (**B**) Representative IF staining of connexin-43 gap junctions (cyan) and cytoskeleton (vimentin, magenta) for confluent MRL/MpJ cells. Scale bar, 100 μm. (**C-F**) Quantification of cell number (**C**), fibronectin deposition (**D**), and cytoskeletal (**E**) and fibronectin alignment (**F**). Female MRL/MpJ cells exhibited significantly greater cell number but diminished ECM deposition and cell-ECM alignment compared to that of male cells. (**G**) Quantification of intercellular communication via connexin-43 expression and (**H**) cellular organization. MRL/MpJ cells displayed comparable gap junction formation and nuclear alignment that were greater than that of male B6 cells. *\*P* < 0.05, *\*\*P* < 0.01, *\*\*\*P* < 0.001, \*\*\*\**P* < 0.0001.

Contrary to our 2D sex differences data, female MRL/MpJ cells displayed significantly lower fibronectin deposition (1.48-fold lower; *P* < 0.0001) and cytoskeletal (*P* = 0.0001) and ECM alignment (*P* = 0.0042) than male MRL/MpJ cells (**Fig. 6C-F**). Female MRL/MpJ cell number remained significantly greater than male MRL/MpJ cells (52.9% higher; *P* < 0.0001) consistent with our 2D data. There was no difference in connexin-43 expression between sexes. Lastly, further supporting our hypothesis, MRL/MpJ cells of both sexes showed significantly greater cytoskeletal and ECM alignment, connexin-43 expression, and nuclear alignment compared to male B6 cells (**Fig. 6H**).

## 4. Discussion

We have previously shown that the improved healing capacity of male MRL/MpJ tendon cells is characterized by increased cellular protrusions, cytoskeletal and cell-deposited ECM alignment, intercellular communication, and proliferation compared to male B6 cells under 2D culture *in vitro*.^23^,^27^ However, given the temporally-distinct sexually dimorphic regulation of this scarless healing capacity in MRL/MpJ tendons^24^, we sought to determine the extent to which these enhanced MRL/MpJ biological processes are preserved between sexes, and therefore constitute a universal regenerative cellular phenotype, or can be leveraged to identify converging pathways that lead to scarless tendon healing. To this end, we comparatively evaluated sex differences in MRL/MpJ cell behavior by modulating topographical and ECM-associated biochemical cues, thereby allowing us to recapitulate key features of the early tendon wound microenvironment.

Our data demonstrated that MRL/MpJ cells exhibited minimal cellular senescence, as evidenced by relatively low ß-gal staining. Under 2D culture, female cells showed significantly lower spreading area, in addition to greater proliferative activity under single-cell and confluent cell densities accompanied by fibronectin-coated substrates or a higher/comparable total amount of cell-deposited fibronectin compared to male cells. Female cells also displayed a lower migratory capacity, greater ECM alignment, and fibronectin-dependent enhanced mechanosensitivity (cellular protrusions, focal adhesion formation, and YAP nuclear localization) compared to male cells. Our previous *in vivo* data revealed decreased activity of MMP-2, an enzyme that degrades fibronectin whose production is regulated by estrogen signaling^30,31^, in female MRL/MpJ tendons compared to male MRL/MpJ tendons during the proliferative phase of wound healing. Jones *et al*.^32^ recently reported a mechanoregulatory mechanism wherein elevated YAP/TAZ signaling suppresses catabolic gene expression that corroborates our *in vitro* and *in vivo* findings. Furthermore, our group and others have postulated a role for cellular protrusion dynamics in sensing and responding to changes in mechanical loading that subsequently drives cytoskeletal and ECM reorganization^23,33^. Taken together, our data suggest a positive feedback mechanism wherein the interaction of female MRL/MpJ cells with wound healing-associated biochemical cues, namely fibronectin, results in an expedited cellular activation response compared to male MRL/MpJ cells.

When comparing MRL/MpJ cell behavior between 2D and 3D culture conditions, sex differences remained for cell spreading area and proliferation. Unexpectedly, female MRL/MpJ cells displayed lower number of cellular protrusions and cellular protrusion formation than male MRL/MpJ cell under 3D culture. Moreover, contrary to our previous 2D findings, male B6 cells displayed comparable protrusion characteristics and greater YAP nuclear localization under 3D culture. However, MRL/MpJ cells of both sexes exhibited significantly greater aspect ratios, a robust indicator of physiological cell elongation, and enhanced cytoskeletal and cell-deposited ECM alignment, ECM deposition, and connexin-43 gap junction expression compared to male B6 cells under 3D culture. Accordingly, these discrepancies in cell morphology suggest that cellular protrusion dynamics may be limited by physical constraints (e.g., fiber diameter, alignment, and porosity and decreased cell spreading). Relatedly, other studies have similarly reported decreased YAP nuclear localization on 3D substrates with higher stiffnesses.^34,35^ Supporting our conflicting 2D and 3D YAP data, the present study employed commercial 3D nanofiber scaffolds composed of PCL, a widely-used material in electrospinning that possesses a tensile modulus close to that of native tendon mechanical properties.

Contrary to our 2D findings, female MRL/MpJ cells showed significantly lower cytoskeletal and cell-deposited ECM alignment than male MRL/MpJ cells. However, a limitation of this study is that these scaffolds resisted cell-mediated fiber recruitment and deformation that would otherwise occur during tendon healing *in vivo*. Consequently, this could have contributed to female MRL/MpJ cells undergoing an altered cellular response and/or disruptions to cell-ECM remodeling concurrent with suppressed ECM catabolism and biophysical overcrowding effects from extensive contact inhibition due to increased cellularity. Interestingly, our data indicate that male MRL/MpJ cells are more capable of overcoming persistent structural disorganization through changes in cell-ECM and/or cell-cell adhesion that ultimately lead to more effective healing outcomes.

Studies investigating fundamental tendon cell biology have been hindered by the notorious phenotypic drift (i.e., loss of tenogenic morphology, markers, and gene expression) observed in primary tendon cell lines.^36–38^ Hussien *et al*.^39^ reported that human tendon-derived stromal cells undergo a dramatic phenotypic drift on conventional tissue-culture plastic (TCP) substrates compared to transcriptional activation of tendon-specific signatures on stiffer substrates. Other efforts to maintain tendon cell behavior have focused on standardizing primary cell isolation methods^40^, applying controlled mechanical stimulation using bioreactor designs^41–44^, and incorporating biochemical cues through exogenous growth factor supplementation and hypoxic culture^45,46^. When considering 3D culture approaches, synthetic scaffolds offer an attractive approach over cell-encapsulated collagen and/or fibrin constructs given their capacity for surface functionalization using ECM proteins and more precise control over the fiber architecture to direct cell attachment, growth, differentiation.^47–49^ Surprisingly, our results revealed that differences between B6 and MRL/MpJ cell processes are largely preserved across 2D and 3D culture environments, which indicates that the complexity of 3D culture platforms may not always be necessary for examining *in vitro* tendon cell behavior despite divergent transcriptional programs compared to 2D substrates. Future work could focus on comprehensively intrinsic phenotypic differences or the cell populations underlying these phenomena to better inform potential mechanistic targets.

In summary, we have shown that MRL/MpJ tendon cells illustrate sex differences in morphology, proliferation, mechanosensitivity, and cell-ECM remodeling *in vitro* that primarily manifest in response to wound healing-associated biochemical cues. We established the utility and advantages of employing 2D and 3D culture platforms with varying topographical cues that will be instrumental in elucidating the mechanistic drivers of regenerative tendon cell biology. Despite these early deviations in cell behavior, we determined that male and female MRL/MpJ cells outperform B6 cells in functional processes associated with regenerative healing outcomes. Collectively, our findings suggest that these regenerative processes are universal to the MRL/MpJ cellular phenotype, which serves as a starting point for identifying shared biological signaling pathways for guiding future therapeutic augmentation strategies that can be employed for a range of tendon healing environments and injury states.

## Acknowledgments

This work was supported by the National Institutes of Health (NIH) under the following award numbers: R01AR608301 (to N.A.P.), R21AR074602 (to N.A.P.), R56AR077239 (to N.A.P.), and S10RR025502 (to Cornell Biotechnology Resource Center). The authors also acknowledge support from the National Science Foundation (NSF) Graduate Research Fellowship Program (GRFP) DGE-1650441 (to J.C.M.) and the Cornell Provost Diversity Fellowship (to J.C.M.). The authors thank Rebecca Bell for insightful discussions in interpreting the study results and Nina S. Tang and Devina Purmessur for helpful technical discussions in using the nanofiber scaffolds.

## Conflicts of Interests

The authors declare that there are no conflicts of interests.

## Author Contributions

JCM conceived the study, designed the experiments, developed protocols for imaging cell-seeded scaffolds, and wrote the manuscript. NAP advised study conception, data interpretation, and writing of the manuscript. JCM, MEB, and MLSP contributed to data collection, analysis, interpretation, and revision of the manuscript.

## Notes

### Competing Interest Statement

The authors have declared no competing interest.

