## Supplemental Figure 1 for "Regenerative MRL/MpJ Tendon Cells Exhibit Sex Differences in Morphology, Proliferation, Mechanosensitivity, and Cell-Matrix Remodeling"

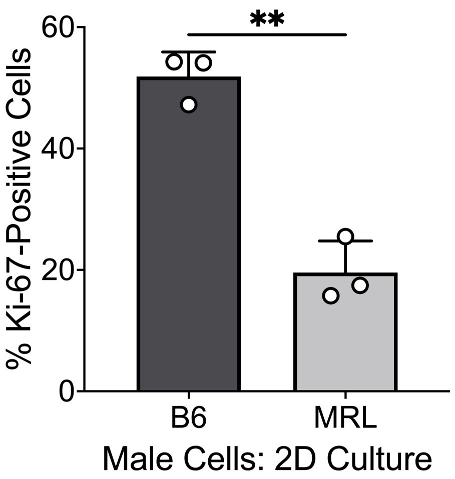


**Supplemental Figure 1. Comparison of 2D cell proliferation between male B6 and MRL/MpJ tendon cells.** Quantification of the percentage of Ki-67-positive cells revealed significantly more proliferating B6 cells than MRL/MpJ cells cultured on TCP substrates. Data is presented as mean ± SEM. $^{**}P < 0.01$.
